# Integration of Kinetic Data into Affinity-Driven Models for Improved T Cell-Antigen Specificity Prediction

**DOI:** 10.1101/2024.06.17.599469

**Authors:** Zahra S. Ghoreyshi, Hamid Teimouri, Anatoly B. Kolomeisky, Jason T. George

## Abstract

T cell receptor (TCR) and peptide-major histocompatibility complex (pMHC) interactions that result in T cell activation are complex and have been distinguished by their equilibrium affinity and kinetic profiles. While prior affinity-based models can successfully predict meaningful TCR-pMHC interactions in many cases, they occasionally fail at identifying TCR-pMHC interactions with low binding affinity. This study analyzes TCR-pMHC systems for which empirical kinetic and affinity data exist and prior affinity-based predictions fail. We identify a criteria for TCR-pMHC systems with available kinetic information where the introduction of a correction factor improves energy-based model predictions. This kinetic correction factor offers a means to refine existing models with additional data and offers molecular insights to help reconcile previously conflicting reports concerning the influence of TCR-pMHC binding kinetics and affinity on T cell activation.

## Introduction

The adaptive immune response relies on T cell activation upon encountering antigenic peptide sequences. T cell specificity stems from the T cell receptor (TCR)’s selective recognition of antigenic peptides presented on the cell surface via major histocompatibility complex (MHC) molecules (1). The human T cell repertoire is comprised of a huge number (*∼* 10^8^) (2) unique TCRs, which collectively confer broad immunity against a variety of antigenic peptides presented on MHC (pMHC). Central thymic selection and peripheral tolerance mechanisms train T cells to distinguish self from non-self signatures (3, 4) and result in a TCR repertoire having variable specificities directed against a particular set of non-self antigens.

Reliable prediction of TCR specificity against antigens of interest remains an active area of research with broad implications for a better more microscopic understanding of immune responses during infection, autoimmunity, and cancer. This problem is challenging because all available training data are sparse in the immense sequence space of TCR and antigen pairs. Several inferential modeling strategies have been proposed, including those based on TCR-peptide primary amino acid sequences (5–11). As an alternative, recently developed random energy models provided a theory-driven approach to understanding repertoire-level T cell tolerance and antigen recognition (12–15). These affinity-based models have subsequently led to the development of inferential biophysical approaches, which leverage known crystal structures to predict TCR-pMHC interactions based on pairwise amino acid potentials trained using available affinity (*K*_d_) data on TCR-pMHC systems (16–18).

The above biophysical models perform well relative to sequence-based models when evaluated using distinct unseen TCRs during training. Incorporating structural information permits reliable predictions using a small fraction of the data needed to train alternative models. Despite these significant advantages, affinity-based modeling still poorly explains some systems, and the molecular reasons for these observations are not well understood. TCR-pMHC interaction kinetics seem to be important in eliciting T cell responses (19, 20), but, relative to affinity measurements, there is a very limited amount of kinetic data (e.g., association and dissociation rate constants *k*_on_ and *k*_off_) available for previously studied human TCR-pMHC systems. In some situations, more information is available on the kinetic responses of TCR-pMHC interactions. The observation of decreased performance of the affinity-based model in certain cases led us to investigate whether incorporating relevant kinetic information, where available, could improve the model predictions.

Here, we studied whether the relatively weaker performance in certain cases could be improved by the inclusion of kinetic information, such as effective TCR-pMHC association (*k*_on_) and dissociation (*k*_off_) rates. To analyze this, we developed a simple method to directly integrate kinetic data, where available, into our previously developed affinity-based models. This approach employs a correction factor derived from kinetic arguments to adjust the predicted affinity values, aiming to more accurately reflect the true binding dynamics observed in physiological conditions. We apply our integrated method to evaluate HLA-A*02-restricted 1G4 TCR binding to mutations of NY-ESO-1 peptide ligands. Our findings demonstrate that the inclusion of the kinetic parameter *k*_on_ is required to resolve TCR specificity while considering the thermodynamic *K*_d_ alone is insufficient. Moreover, the incorporation of *k*_on_ significantly improves the prediction accuracy in our affinity-driven model, which previously was ineffective at resolving NY-ESO-1-specific TCRs. This approach offers a method by which complementing affinitydriven modeling with kinetic information can dramatically enhance the quantitative description and our understanding of complex processes in immune response.

## Methods

### TCR-pMHC Binding Dynamics

A T cell receptor with concentration [*TCR*] binds a peptide-bound major histocompatibility species that have concentration [*pMHC*], to create a TCR-pMHC complex with concentration [*TCR* : *pMHC*]. This process can be described as a reversible chemical reaction,

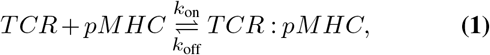

for which the equilibrium dissociation constant of this process (*K*_d_) can be expressed as the ratio of corresponding equilibrium concentrations or the ratio of the association and dissociation rate constants,

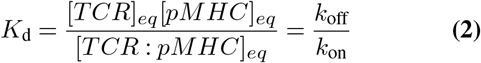

The equilibrium constant *K*_d_ provides a static thermodynamic view of the binding strength while 1*/k*_off_ can be associated with the duration of time that the single TCR-pMHC complex remains intact before disassembly, which might be relevant for T cell signaling response. Therefore, the experimental values of *K*_d_, *k*_off_, and *k*_on_ that are used as input in our model are directly applied to evaluate the efficacy of different TCR-pMHC interactions, guiding the development of our computational models and the interpretation of their outputs.

#### Predictive Structural Affinity Model

To accurately estimate the binding strength of a specific TCR-pMHC pair, we utilized our previously developed RACER pairwise energy framework (16, 17). RACER calculates binding energies by integrating high-throughput data on previously confirmed TCR-peptide interactions and crystal structures to train a residue-specific energy matrix. The energy matrix and available structural templates are then used to quantify TCR-peptide binding. To accomplish this, RACER evaluates the binding interface between peptide antigen and variable (CDR3*α*/*β*) regions via:

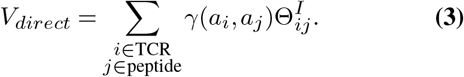

In Eq. 3, *γ*(*a*_*i*_, *a*_*j*_) denotes the pairwise interaction strength between amino acids *a*_*i*_ and *a*_*j*_ at positions *i* and *j* within the indexed TCR and peptide, respectively. The entries of *γ* represent all symmetric pairwise amino acid energies, which are optimized during model training (21). Each interaction is weighted as a sigmoidal function Θ_*ij*_ of the pairwise distances between residues:

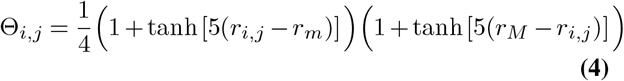

where *r*_*m*_ = 6.5 Å (respectively *r*_*M*_ = 8.5 Å) define the minimum (respectively maximum) distances over which the interaction remains significant.

RACER computes the energy of test TCR-pMHC pairs (*E*_test_). These values are compared against energy values for an ensemble of randomized weak binding pairs (*E*_weak_) binding TCR-pMHC pairs, and the free energy is defined relative to the mean of weak binders: Δ*G ∼ E*_test_ − ⟨*E*_weak_⟩. Experimentally obtained *K*_d_ and calculated approximated free energy changes Δ*G* are assumed to be related via

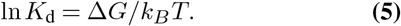

To compare standardized free energy across distinct TCR-pMHC pairs and structural templates, the *z*-score is introduced as a normalized free energy parameter, defined as follows:

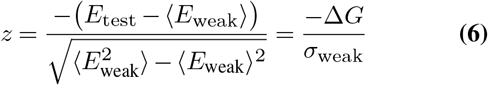

### Integrating Kinetic Data into the Affinity Model

To incorporate additional experimental kinetic data into the RACER framework, we modify the imputed *K*_d_ values. As one can see from Eq. 5, the parameters *K*_d_ and *k*_off_ are expected to vary proportionally across TCR-pMHC pairs that have comparable values for *k*_on_. Previous work has however noted that the variations in on-rate *k*_on_ can significantly influence overall interaction dynamics in the TCR-pMHC binding case(22). To account for this, we introduce a *kinetic correction term, η*_*i*_, to modify the *K*_d_ values for TCR-pMHC pair *i* in cases where on-rate (*k*_on,i_) is available according to the following rule:

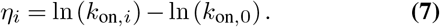

In the above equation, cases where *k*_on,*i*_ exhibits significant deviation away from typical values of *k*_on,0_ of observed systems, *η* can be used to create an ‘effective dissociation equilibrium constant’, 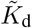, according to:

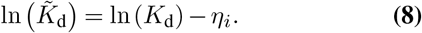

The application of Eq. 7 relies on an empirical analysis of experimentally obtained *k*_on_ values for relevant cases. Our analysis (see the Results Section) ultimately leads us to select a lower limit for *k*_on_ ⋍ 10^5^ below which the correction *η* is subsequently applied.

### Data Acquisition and Analysis

Affinity data for 131 TCR-pMHC systems were sourced from the ATLAS database (23). All available crystal structures for HLA A*02-restricted systems were obtained via the RCSB Protein Data Bank (24). Additional kinetic data were obtained from a recent analysis on the NY-ESO-related peptides’ interaction with the 1G4 TCR (22). This augmented set provided a total of 11 additional TCR-pMHC pairs (Table S1 and Table S2).

## Results

Fig. 1 presents an overview of our approach, which involves integrating kinetic data in the form of a corrected affinity score. Our primary goal is motivated by the question of whether or not the incorporation of additional kinetic information into a structural model trained on affinity data could improve the predictions, particularly in cases where available affinity data are insufficient for explaining relevant TCR-pMHC interactions.

**Fig. 1.**
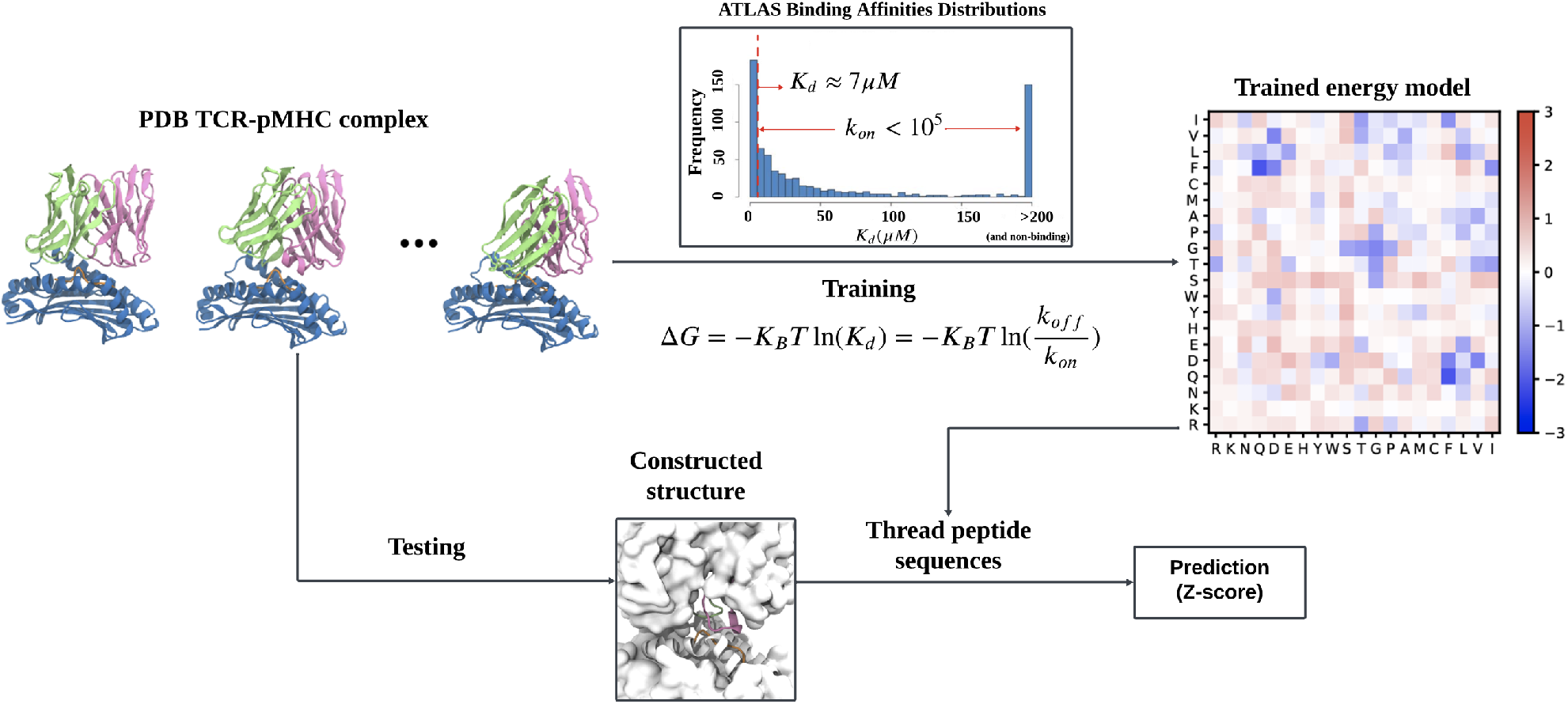
Schematic depiction of the training and testing phases within RACER, illustrating the impact of the *k*_on_ binding parameter on the trained energy model. For each TCR-pMHC pair with *K*_d_ *>* 7*µM* and *k*_on_ *<* 10^5^, a correction factor was applied.

### 1G4-NY-ESO as a Candidate TCR-pMHC System

The NY-ESO peptide and corresponding 1G4 TCR variants served as appropriate candidates as sufficient structural information exists for this case, yet despite this, RACER-m, a multi-template approach within the RACER framework, was unable to correctly assess 50% of available cases even when they were explicitly included in training (17). An additional independent dataset generated affinity and kinetic data in the form of *k*_on_, *k*_off_, and *K*_d_ for additional 1G4-NY-ESO (TCR-pMHC) mutational variants (22).

Plotting the effective off-rate *k*_off_ as a function of the equilibrium constant *K*_d_ for NY-ESO-associated cases demonstrated a statistically significant direct correlation between affinity and kinetic parameters, and expected inverse trends were also found for respectively *k*_on_ vs. *K*_d_ (Fig. 2a-b). Significant inverse correlation was also observed between *k*_on_ and *k*_off_ (Fig. 2c). However, this behavior was inordinately influenced by a minority of cases with large *k*_on_. Removal of the three NY-ESO-mutant TCR-peptide pairs having lowest *k*_on_ values was enough to break the observed correlation between *k*_off_ and *K*_d_ (Fig.2d, Table S3).

**Fig. 2.**
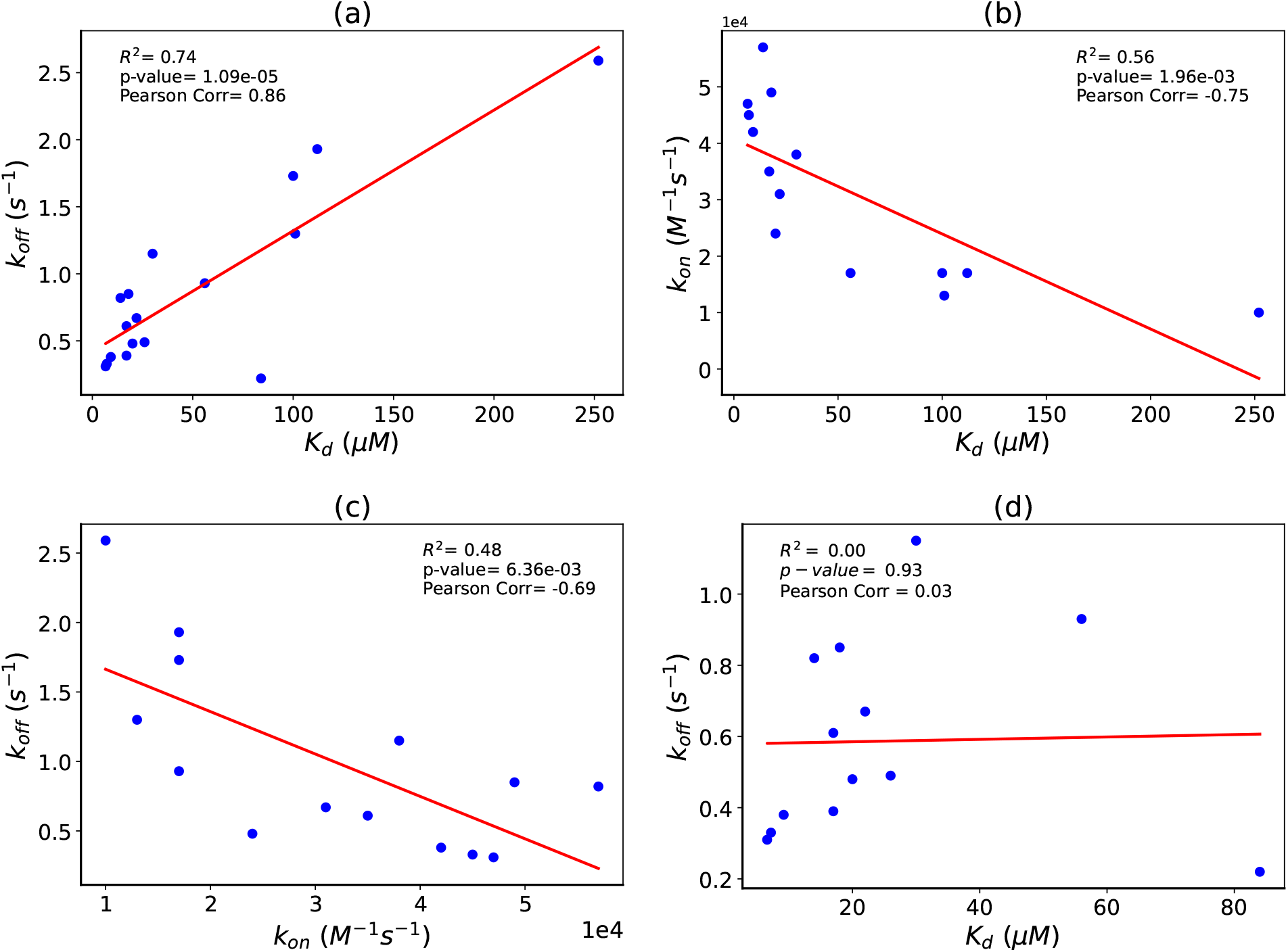
Correlation Analyses of TCR-pMHC Kinetic and Affinity Parameters for NY-ESO Cases. This figure explores the interrelations between kinetic and affinity parameters across all NY-ESO cases. (a) Demonstrates the correlation between dissociation constant *K*_d_ and dissociation rate *k*_off_, illustrating their mutual dependencies. (b) Examines the relationship between *K*_d_ and association rate *k*_on_, assessing how association dynamics influence binding affinity. (c) Analyzes the interplay between *k*_off_ and *k*_on_, highlighting the balance of association and dissociation in binding kinetics. (d) The correlation between *K*_d_ and *k*_off_ specifically for a subset of NY-ESO cases.

Because the RACER-m model attempts to resolve meaningful TCR-peptide pairs based on an affinity threshold (defined to be *K*_d_ *≈* 7*µM*), we hypothesized that cases with predicted *K*_d_ values exceeding this threshold, especially those with the smallest *k*_*on*_ values, would benefit significantly from a kinetic normalization of the affinity model’s value. In the regime of low *k*_on_, small fluctuations can have a large impact on the relationship between *K*_d_ and *k*_off_. This supports the need for the kinetic correction term, which is further discussed in the Results Section.

### Selection of a kinetic correction term recovers identification of favorable interaction pairs

We applied a similar analysis on the subset of the ATLAS dataset for which TCR-pMHC systems had an affinity and kinetic data (Fig.3). Again, significant correlation have been found in relations between *k*_off_ vs. *K*_d_ and the removal of cases with low *k*_on_ disrupts the observed correlation between *k*_off_ and *K*_d_ (Fig. 3a-b). For these cases, when *k*_on_ *>* 10^5^, the values of *K*_d_ and *k*_off_ are concentrated in a narrow range. Conversely, when *k*_on_ *<* 10^5^, *K*_off_ and *K*_*d*_ are highly variable (Fig. 3c-d). While this behavior appears to affect NY-ESO cases most strongly, additional TCR-pMHC systems from distinct functional clusters also exhibit significant variation in *k*_off_ and *K*_d_, including TCRs specific for GILGFVFTL and ILAKFLHWL peptides. These findings are consistent with another report analyzing the 1G4-NY-ESO TCR-peptide, wherein the authors utilized a slightly different approach to account for widely variable *K*_on_ values (22).

**Fig. 3.**
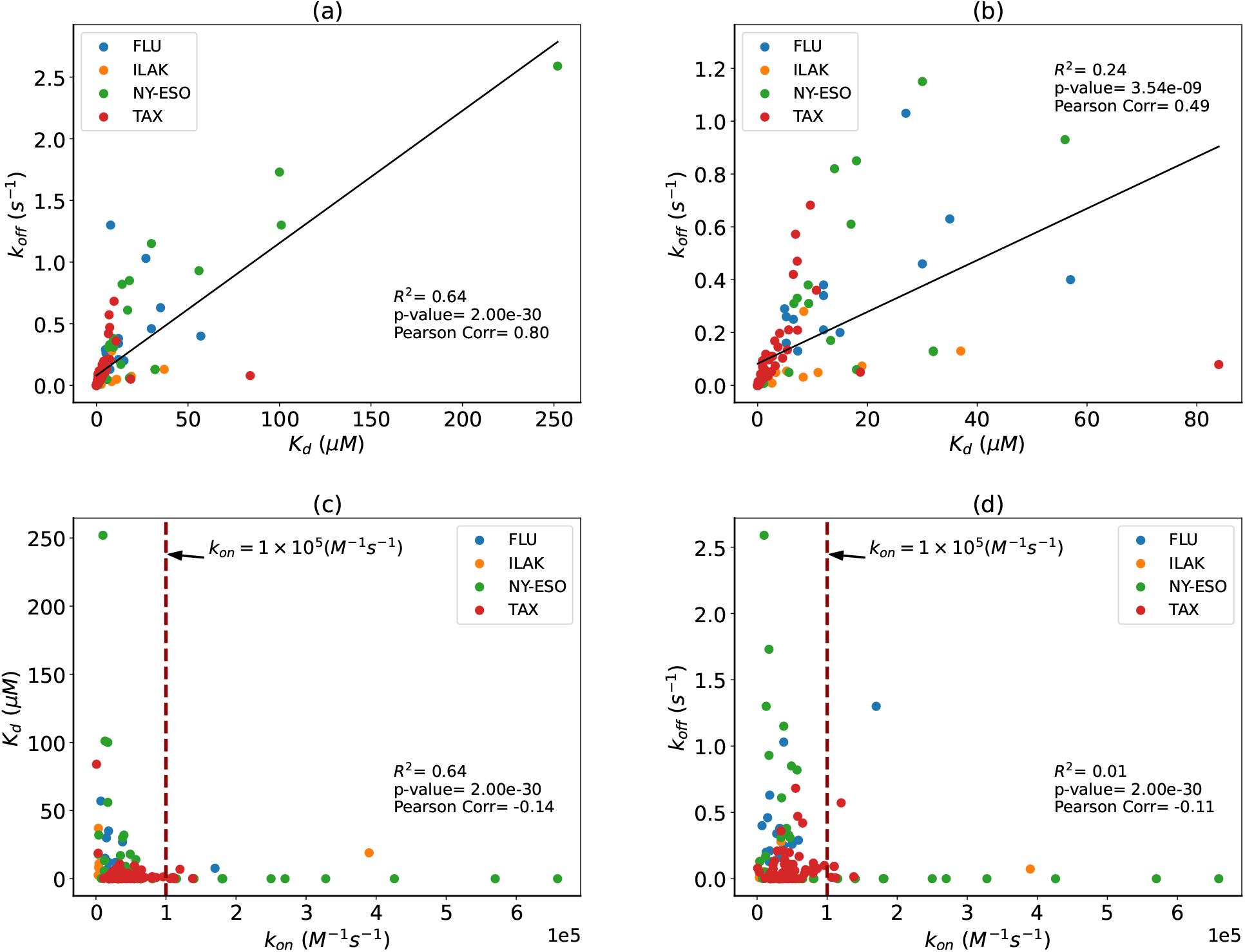
Analysis of Model Fitting to Data Subsets with Larger Variability in *k*_on_. (a) displays the correlation between *K*_d_ and *k*_off_ for the entire ATLAS dataset. (b) focuses on a subset of the ATLAS dataset, excluding cases with high variability in *k*_on_. These panels collectively highlight how different rates of *k*_on_ influence the model’s ability to fit the data, emphasizing the impact of *k*_on_ on binding parameter correlations. (c) illustrates the distribution of *k*_on_ versus *K*_d_, and (d) shows *k*_on_ versus *k*_off_ for all cases in the ATLAS dataset.

Given the above findings, we now introduce a correction fac-tor based on kinetic normalization into our affinity estimates (Eq. 8) for systems having low *k*_on_ (Eq. 7). The correction term effectively incorporates kinetic information in the regime of *k*_on_ *<* 10^5^, for which the explicit incorporation of observable *k*_off_ is expected to reflect the true binding dynamics more accurately.

To evaluate whether this correction term can successfully distinguish meaningful TCR-pMHC pairs, we incorporate this adjustment into the discrimination step based on standardized z-score (Eq. 6). The predicted z-scores as a function of log(*K*_d_) are plotted both before (Fig. 4a) and following (Fig. 4b) correction for NY-ESO-specific TCRs. Using the previously established z-score cutoff, our correction demonstrates effectively recognized NY-ESO-specific TCRs, most of which would have been unidentified. This simple correction results in over 90% of cases being correctly identified. Table 1 and Table 2 display a summary of each case’s corrections.

**Table 1.** Z-scores and Δ*G* values for 1G4 TCR with NY-ESO peptide and its mutations before and after applying the correction factor.

| Peptide name | $k_{on} \times 10^3 (M^{-1}s^{-1})$ | $\eta_i$ | $Z - score$ | $\tilde{Z} - score$ | $\Delta G (kcal mol^{-1})$ | $\tilde{\Delta G} (kcal mol^{-1})$ |
| --- | --- | --- | --- | --- | --- | --- |
| ESO-9C | 57 | 0.63 | 1.1706 | 2.8172 | -2.6642 | -4.2259 |
| ESO-9L | 17 | 0.58 | 0.8390 | 2.6618 | -2.5551 | -3.9928 |
| ESO-9V | 45 | 0.39 | 1.0693 | 2.2976 | -2.4801 | -3.4468 |
| ESO-9A | 47 | 0.44 | 1.0598 | 2.4064 | -2.5189 | -3.6096 |
| ESO-3I | 35 | 0.14 | 0.9459 | 1.9619 | -2.5145 | -2.9429 |
| ESO-3M | 42 | 0.36 | 1.061 | 2.3252 | -2.5959 | -3.4878 |
| ESO-3Y | 38 | 0.23 | 0.9250 | 1.9042 | -2.2861 | -2.8562 |
| ESO-4D | 10 | 1.11 | 0.9358 | 2.0054 | -2.4172 | -5.1687 |
| ESO-6V | 49 | 0.48 | 0.9615 | 2.4148 | -2.4324 | -3.6222 |
| ESO-6T | 13 | 0.85 | 0.9548 | 1.7045 | -2.2861 | -4.3931 |
| ESO-7H | 17 | 0.58 | 0.5530 | 1.0649 | -1.3070 | -2.7447 |

**Table 2.** Z-scores and Δ*G* values for 1G4 TCR and its mutations before and after applying the correction factor.

| TCR name | $k_{on} \times 10^3 (M^{-1}s^{-1})$ | $\eta_i$ | $Z - score$ | $\tilde{Z} - score$ | $\Delta G (kcal mol^{-1})$ | $\tilde{\Delta G} (kcal mol^{-1})$ |
| --- | --- | --- | --- | --- | --- | --- |
| 1G4-mu1 | 17.8 | 0.16 | 0.3758 | 2.3714 | -0.9691 | -6.6050 |
| 1G4-mu2 | 34 | 0.12 | 0.6167 | 2.6135 | -0.9633 | -5.2277 |
| 1G4-mu3 | 4 | 2.03 | 0.5893 | 3.9464 | -2.0482 | -7.8929 |
| 1G4-mu4 | 11 | 1.01 | 1.0706 | 3.0421 | -2.9691 | -6.0842 |

**Fig. 4.**
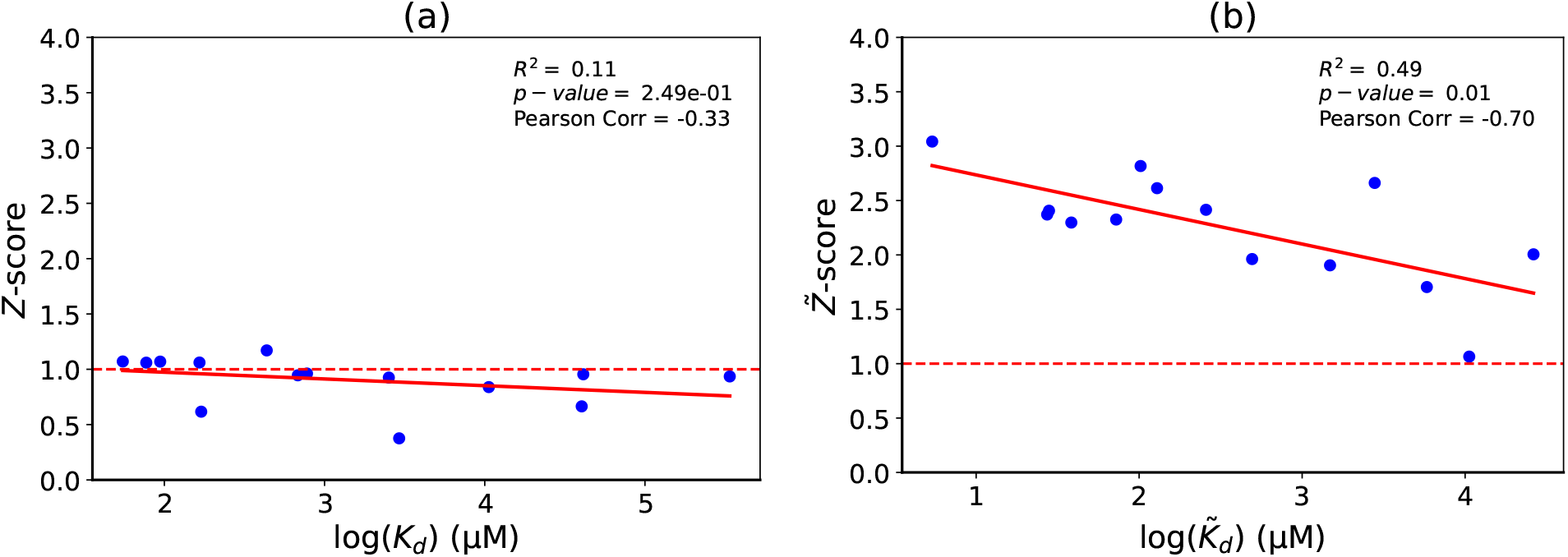
Detailed Analysis of TCR-pMHC Binding Parameters for Select NY-ESO Cases. Comparison of RACER z-scores and experimental *K*_d_ values for NY-ESO cases with high *K*_d_ values, providing insight into the model’s effectiveness in predicting TCR-pMHC interactions. (a) Z-scores before applying the correction factor. (b) Z-scores after applying the correction factor, illustrating the improved predictions. This comparison highlights the precision of the RACER model in handling cases with significant variability, thereby illustrating its utility in refining predictions for complex immune responses.

### Incorporation of kinetic correction term leads to improved model predictions for meaningful TCR-pMHC interactions

The kinetic correction factor can improve the predictions on NY-ESO-specific TCRs. In the next step, we implement this correction into the existing RACER approach applied to distinct TCR-pMHC pairs with experimentally confirmed thermodynamic parameters (23, 25–28). To assess this, we applied the optimized RACER model to predict TCR-pMHC interactions, restricting our analysis to TCR-pMHC systems in ATLAS with available kinetic data. Our results are reported in Fig. 5 and compare RACER predictions before and after accounting for the correction. Cases for which RACER had previously predicted strong binding did not observe significant changes in the predicted binding energy value. This was most evident in the GILGFVFTL (FLU) antigen-specific TCRs system. Such cases also tended to correspond to TCR-pMHC pairs where the correction criterion was not satisfied (Fig. 5 gray values). This contrasted starkly with the remaining cases, most of which RACER either predicted as a borderline favorable interaction or failed outright. Our kinetic correction successfully improved the original affinity-based predictions in these cases, representing experimentally confirmed favorable TCR-pMHC systems. Lastly, to observe the effects of available kinetic corrections as a function of *K*_d_, we plotted pre-normalization and post-normalization binding energies for all cases with kinetic data rank-ordered by the original *K*_d_ values (Fig. S1). As expected, we find smaller pre-normalization vs. post-normalization case changes as *K*_d_ values decrease.

**Fig. 5.**
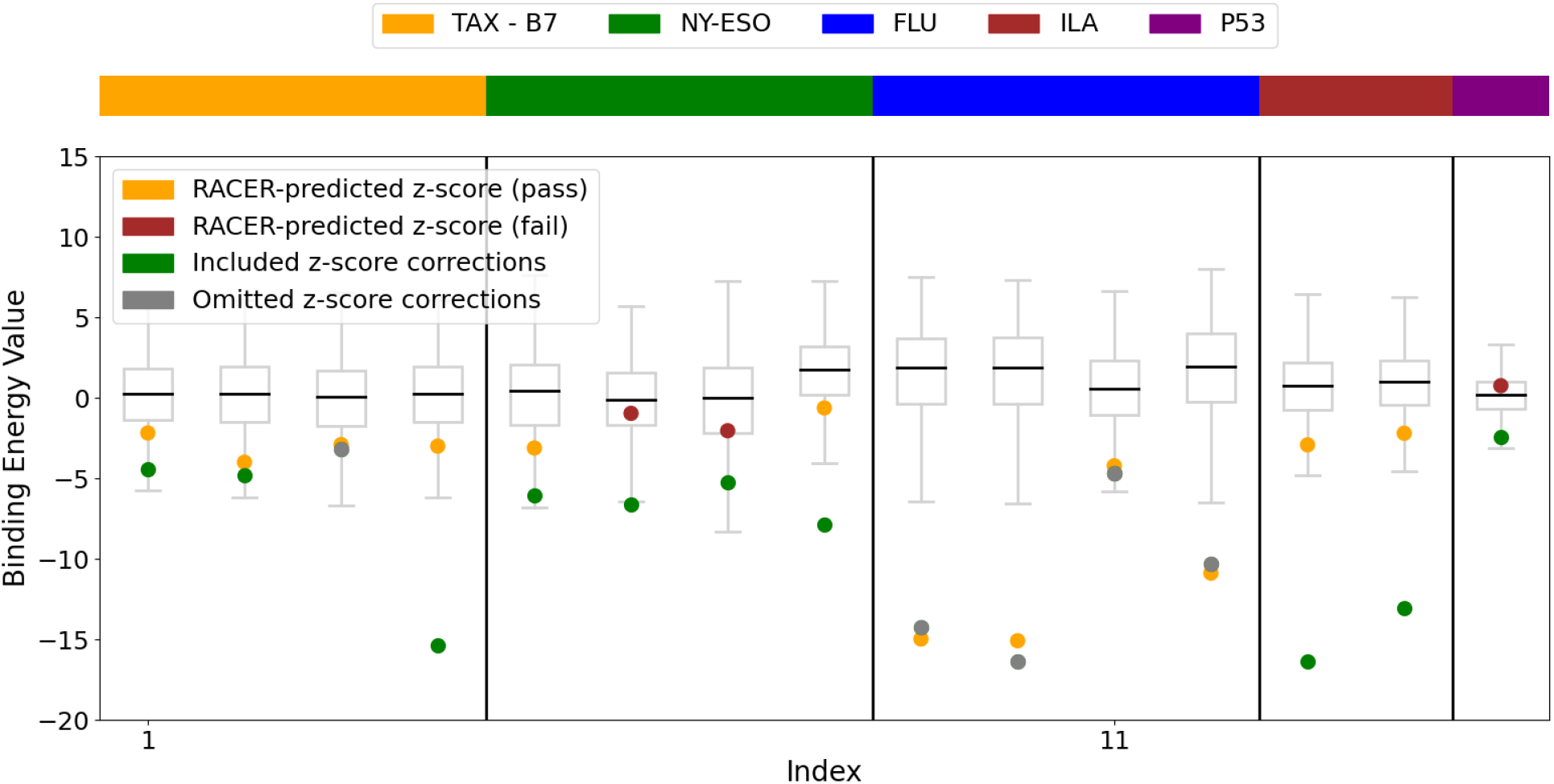
Performance on ATLAS dataset before and after applying the correction factor. The box plots illustrate the distribution of binding energy values, highlighting the variability. The correction factor improves prediction accuracy, particularly in the higher affinity interactions (represented by the orange dots), which is consistent with the lower *K*_d_ values demonstrating less improvement.

## Discussion

Reliable predictions of TCR specificity are still quite challenging partly because a complete description of TCR-pMHC interactions must account for complex interaction dynamics, downstream signaling, and orchestration of cytokine release. Binding affinity, the most widely available thermodynamic characterization of TCR-pMHC interactions, has been successfully applied to train structural models to achieve TCR-pMHC specificity predictions with reasonable accuracy. Despite this, including additional physical parameters, such as binding kinetics, p-MHC abundance on the cell surface, and T cell costimulatory signals, seems necessary to characterize T cell behavior fully.

Our goal here was to investigate with more detail the TCR-pMHC situations for which predictive models trained strictly on binding affinity data performed poorly. Given their inability to be accurately accounted for using available affinity data, these cases, including NY-ESO and several other TCR-pMHC functional systems, provided an opportunity to incorporate and test meaningful kinetic data into model predictions. Across all publicly available TCR-pMHC systems, a dramatic increase in the variability of *k*_off_ emerged for sufficiently low *k*_on_ values. This observation motivated us to introduce kinetic threshold criteria for refining our earlier modeling approach.

While successful on the limited set of currently available systems, subsequent thermodynamic characterization of additional TCR-pMHC cases will be needed to evaluate whether this empirical cutoff scheme can be applied to immunologically distinct systems. One advantage of our threshold-based approach is that it offers a simple criterion for determining when kinetic corrections are needed. For instance, in several TAX-and FLU-specific cases for which our threshold criterion was not met, utilizing the kinetic corrections yielded minimal deviations from the original z-score predictions. This observation was held for cases with predicted scores both close to and far from the critical cutoff for activation.

We assessed the binding affinity of both wild-type and variant 1G4 TCRs to NY-ESO and NY-ESO-variant peptides. Our dataset reveals that pMHC recognition does not always align well with either *K*_d_ or *k*_off_ for cases having lower *k*_on_. We augmented our analysis to include additional TCR-pMHC systems with considerable variation in *k*_on_ to more comprehensively explore *k*_on_’s impact on pMHC recognition. Our results align with prior studies that found consistent *k*_on_ values were most important in successfully correlating TCR-pMHC recognition with either *K*_d_ or *k*_off_ (29). In many cases however, TCR-pMHC recognition has been found success-fully correlating with either *K*_d_(1, 30, 31) or *k*_off_(32–36).

Our results demonstrate that one can address the practical question of when and how to effectively incorporate variable thermodynamic features into inferential models of TCR-pMHC specificity by identifying and incorporating suitable correction factors. Moreover, such adjustments when properly introduced can successfully encode independent information (off-rate) that is otherwise distinct from the original model biophysical feature (affinity). We remark that while our correction factor is incorporated using thermodynamic theory, the precise criteria for inclusion were made empirically based on all available observations. As a result, our prediction refinements are limited by the availability and accuracy of kinetic data.

Our theoretical method works reasonably well for low values of the on-rates, and one can provide a more microscopic explanation of this observation. The equilibrium constant *K*_*d*_ provides a measure of binding affinity if these events can be measured for a long time. For large on-rates after some time the *K*_*d*_ can be well evaluated. However, when the on-rates are small due to limited time for immune response processes one cannot rely on *K*_*d*_, and the kinetic information provides a better description of the effective binding affinity.

While successfully applied to multiple TCR-pMHC systems, this approach will require further validation on distinct functional systems. These follow-up analyses will benefit from additional TCR-pMHC kinetic information, particularly in cases having low on-rates. The above approach demonstrates promise for broadening the class of encodable information useful to an existing, affinity-based model. In this case, kinetic data was included in a straightforward way using foundational thermodynamic principles. Future research should focus on expanding this approach to include additional features that are commonly measured, such as pMHC abundance and cytokine release, in a biophysically meaningful manner. Such model improvements will broaden the utility of predictive models and in doing so will enhance therapeutic strategies to predict TCR-pMHC interactions with applications to cancer immunology, infectious diseases, and autoimmunity.

## Acknowledgments

JTG was supported by the Cancer Prevention Research Institute of Texas (RR210080). JTG is a CPRIT Scholar in Cancer Research. ABK was supported by the Welch Foundation (C-1559), and by the Center for Theoretical Biological Physics sponsored by the NSF (PHY-2019745).

