## Supplementary Information for "Integration of Kinetic Data into Affinity-Driven Models for Improved T Cell-Antigen Specificity Prediction"

| **Peptide Name** | **Peptide Sequence** | $K_d$ ($\mu M$) | $k_{off}$ ($s^{-1}$) | $k_{on}$ $\times 10^3$ ($M^{-1}s^{-1}$) | $\Delta G$ (kcal mol$^{-1}$) |
|---|---|---|---|---|---|
| ESO-9C | SLLMWITQC | 14 ± 1 | 0.82 ± 0.01 | 57 ± 3 | -6.9 ± 0.0 |
| ESO-9L | SLLMWITQL | 56 ± 6 | 0.93 ± 0.05 | 17 ± 2 | -6.0 ± 0.1 |
| ESO-9V | SLLMWITQV | 7.2 ± 0.5 | 0.33 ± 0.01 | 45 ± 4 | -7.3 ± 0.0 |
| ESO-3A | SLAMWITQV | 6.6 ± 0.5 | 0.31 ± 0.01 | 47 ± 4 | -7.4 ± 0.0 |
| ESO-3I | SLIMWITQV | 17 ± 1 | 0.61 ± 0.04 | 35 ± 3 | -6.8 ± 0.0 |
| ESO-3M | SLMMWITQV | 9.2 ± 0.2 | 0.38 ± 0.01 | 42 ± 1 | -7.1 ± 0.0 |
| ESO-3Y | SLYMWITQV | 30 ± 1 | 1.15 ± 0.04 | 38 ± 1 | -6.4 ± 0.0 |
| ESO-4D | SLLDWITQV | 252 ± 12 | 2.59 ± 0.15 | 10 ± 1 | -5.1 ± 0.0 |
| ESO-6V | SLLMWVTQV | 18 ± 0 | 0.85 ± 0.03 | 49 ± 2 | -6.8 ± 0.0 |
| ESO-6T | SLLMWTTQV | 101 ± 5 | 1.30 ± 0.03 | 13 ± 1 | -5.7 ± 0.0 |
| ESO-7H | SLLMWIHQV | 100 ± 8 | 1.73 ± 0.09 | 17 ± 2 | -5.7 ± 0.0 |

Table S1: Binding affinities, kinetic rates, and thermodynamic data of 1G4 TCR with NY-ESO peptide and its mutations. Data is adapted from [1]

| **TCR Name** | **Peptide Sequence** | $K_d$ ($\mu M$) | $k_{off}$ ($s^{-1}$) | $k_{on}$ $\times 10^3$ ($M^{-1}s^{-1}$) | $\Delta G$ (kcal mol$^{-1}$) |
|---|---|---|---|---|---|
| 1G4-mu1 | SLLMWITQC | 0.0816 ± 0.0 | 0.00145 ± 0.0 | 17.8 ± 1 | -9.67 ± 0.0 |
| 1G4-mu2 | SLLMWITQC | 9.3 ± 0.0 | 0.31 ± 0.0 | 34 ± 1 | -6.86 ± 0.0 |
| 1G4-mu3 | SLLMWITQC | 32 ± 0.0 | 0.13 ± 0.0 | 4 ± 1 | -6.13 ± 0.0 |
| 1G4-mu4 | SLLMWITQV | 5.7 ± 0.0 | 0.049 | 11 ± 2 | -7.15 ± 0.0 |

Table S2: Binding affinities, kinetic rates, and thermodynamic data of 1G4 TCR and its mutations with the NY-ESO peptide. Data is adapted from [2]

| **pMHCs Removed** | $R^2$ | **Pearson-Corr** | **p-value** | **Spearman-Corr** | **p-value** |
|---|---|---|---|---|---|
| 0 | 0.7320 | 0.8556 | 2.3806e-05 | 0.6696 | 4.5474e-03 |
| 1 | 0.3839 | 0.6196 | 1.3751e-02 | 0.5987 | 1.8355e-02 |
| 2 | 0.17 | 0.4166 | 1.3838e-01 | 0.5105 | 6.2172e-02 |
| 3 | 0.0007 | 0.0261 | 9.3264e-01 | 0.3879 | 1.9031e-01 |
| 4 | 0.0032 | 0.0564 | 8.6186e-01 | 0.3818 | 2.2071e-01 |
| 5 | 0.0194 | -0.1393 | 6.8284e-01 | 0.3052 | 3.6137e-01 |
| 6 | 0.0095 | -0.0974 | 7.8888e-01 | 0.3769 | 2.8300e-01 |

Table S3: Examining how models conform to partial datasets

| peptide | TCR | $K_D$ ($\mu$M) | $k_{\mathrm{off}}$ ($s^{-1}$) | $k_{\mathrm{on}}$ ($M^{-1}$ $s^{-1}$) |
|---|---|---|---|---|
| native Tax | B7 | 1.35 ± 0.04 | 0.13 | 9.60 ± 0.03 $\times 10^4$ |
| Tax-Y5F | B7 | 6.49 ± 0.17 | 0.42 | 6.50 ± 0.02 $\times 10^4$ |
| Tax-Y5W | B7 | 35 ± 1 | 0.13 | 3.7 ± 0.1 $\times 10^3$ |
| native TCR | B7 Y104$\beta$A | 2.99 ± 0.10 | 0.13 | 4.2 ± 0.1 $\times 10^4$ |

Table S4: Kinetic and affinity measurements for Wild-Type and mutant B7 recognition of native and position 5 variants of the Tax peptide at 25C. Data for the native and Y5F peptides are adapted from Davis et al. (2005) [3], while data for the Y5W peptide and TCR mutants are sourced from Davis et al. (2007) [4].

| peptide | TCR | $K_D$ ($\mu$M) | $k_{\mathrm{off}}$ ($s^{-1}$) | $k_{\mathrm{on}}$ ($M^{-1}$ $s^{-1}$) |
|---|---|---|---|---|
| native FLU | JM22 | 5.2 | 0.16 | 3.1 $\times 10^4$ |
| native FLU | JM22 S99A | 4.9 | 0.29 | 59 $\times 10^4$ |
| native FLU | JM22 Y101A | 7.7 | 1.3 | 170 $\times 10^4$ |
| native FLU | JM22 Y101F | 12 | 0.21 | 18 $\times 10^4$ |

Table S5: Kinetic and affinity measurements for Wild-Type and mutant JM22 recognition of native and position 99, 100, and 101 variants of the FLU TCR at 37C. Data for the native is adapted from ATLAS dataset [2], while data for the TCR mutants are sourced from Ishizuka et al. (2018) [5].

| Peptide | TCR | $K_D$ ($\mu$M) | $k_{\mathrm{off}}$ ($s^{-1}$) | $k_{\mathrm{on}}$ ($M^{-1}$ $s^{-1}$) |
|---|---|---|---|---|
| p53R175H | 12-6 | 1.3 | 0.032 | 2.5 $\times 10^4$ |

Table S6: Kinetic and affinity measurements for Wild-Type 12-6-P53 recognition of native at 37C. Data is adapted from Wu. et al. (2020) [6].

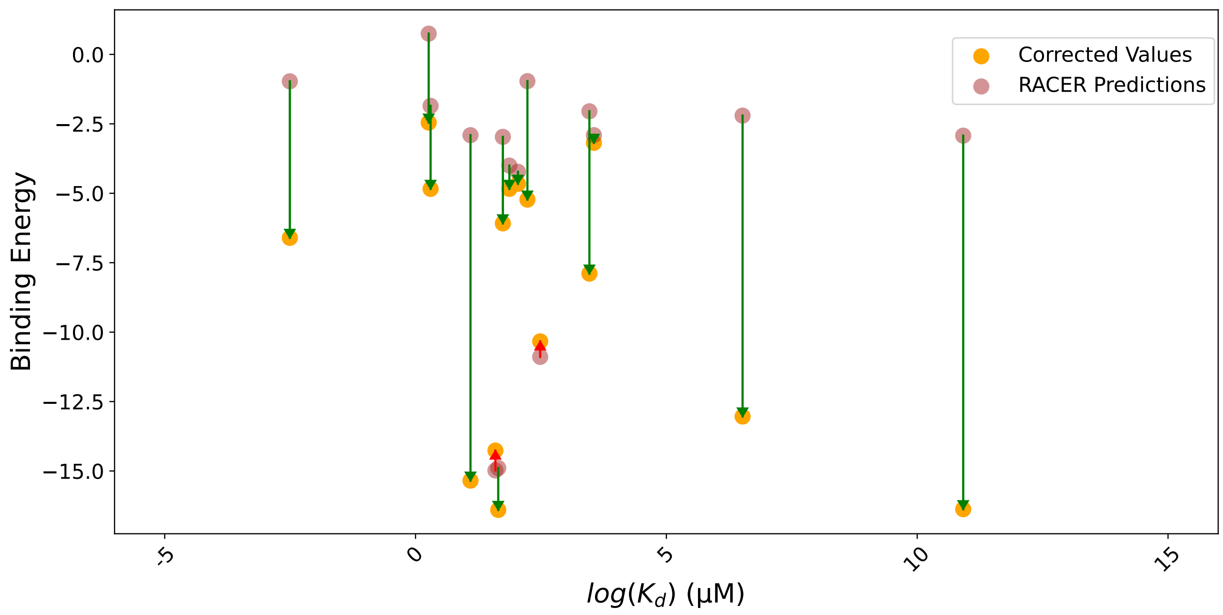


Figure S1: **Binding energy vs. rank of** $\log(K_\mathbf{d})$ **for TCR cases.** The orange dots represent corrected values after applying the correction factor, while the red dots represent the original RACER predictions. The correction factor was applied to improve the accuracy of affinity estimates, particularly at higher affinity interactions. The FLU and TAX cases shows less improvement due to its lower $K_\mathrm{d}$ values compared to other cases.
